# Integrated physiological performance and *Nax1*-mediated sodium exclusion reveal mechanisms of salinity tolerance in spring wheat (*Triticum aestivum* L.)

**DOI:** 10.64898/2026.03.04.709707

**Authors:** Md. Monwar Hossain, Md. Hasanuzzaman, Md. Abul Kalam Azad, Mohammad Nur Alam

**Author notes:** Correspondence to Md. Hasanuzzaman, and Mohammad Nur Alam. (MMH); (MAKA).

## Abstract

Soil salinity is a rapidly intensifying abiotic stress that significantly limits wheat productivity, particularly in coastal and irrigated agroecosystems. Although sodium (Na^+^) ion exclusion has been recognized as a key tolerance mechanism, the integration of physiological performance with *Nax1*-mediated molecular regulation among regionally adapted wheat genotypes remains insufficiently characterized. The present study aimed to dissect salinity tolerance by combining hydroponic phenotyping, multivariate trait analysis, molecular marker profiling, and quantitative expression analysis of the Na^+^ ion transporter gene *Nax1*. Seventeen spring wheat genotypes were evaluated under four salinity levels (0.0, 10, 12, and 14 dS m⁻¹). Germination and survival rate, shoot and root growth, and biomass accumulation were measured. Principal component analysis (PCA) and hierarchical clustering were performed to classify genotypes, while SSR (simple sequence repeat) and *Nax*-linked markers assessed genetic diversity. Relative *Nax1* expression was quantified using qRT-PCR (quantitative real-time polymerase chain reaction). Salinity significantly reduced germination, survival, elongation, and biomass, with strong genotype-dependent variation. Multivariate analyses clearly separated tolerant and sensitive genotypes, with biomass retention and survival contributing most to total variation. Marker analysis revealed moderate genetic polymorphism. Notably, tolerant genotypes exhibited 3-6-fold induction of *Nax1* under severe salinity, positively correlating with biomass maintenance. These findings demonstrate that salinity tolerance in wheat is associated with coordinated physiological resilience and enhanced *Nax1*-mediated Na⁺ ion exclusion, thereby advancing mechanistic understanding and supporting molecular-assisted breeding for salt-affected environments.

## Introduction

Soil salinity is a major abiotic stress limiting crop productivity globally [1-5]. These studies emphasize osmotic stress, ionic toxicity, and disruption of cellular homeostasis as central constraints to plant performance under saline conditions. Soil salinity is one of the most severe abiotic stresses limiting agricultural productivity worldwide. More than 800 million hectares of land are affected by salinity, and nearly 20-30% of irrigated agricultural land is estimated to be salt-affected, with projections indicating continued expansion due to climate change, sea-level rise, and inappropriate irrigation practices [6, 7]. Wheat, a staple cereal providing approximately 20% of global caloric intake, is moderately sensitive to salinity, particularly during germination and early vegetative growth [8, 9]. As global food demand continues to rise, improving wheat resilience to saline environments has become a critical priority for sustainable crop production. Salinity imposes two major phases of stress on plants: an initial osmotic phase that restricts water uptake and a subsequent ionic phase characterized by toxic accumulation of Na⁺ and Cl⁻ in plant tissues [10, 11]. Osmotic stress rapidly reduces cell expansion and shoots growth, while prolonged exposure leads to ionic imbalance, membrane destabilization, and oxidative damage [12]. Excess Na⁺ competes with K⁺ at critical binding sites, disrupting enzymatic processes and impairing photosynthetic efficiency [13, 14]. Consequently, salinity significantly reduces germination percentage, seedling survival, biomass accumulation, and ultimately grain yield.

Plants have evolved multiple adaptive strategies to cope with salinity, including osmotic adjustment, ion compartmentalization, antioxidant defense activation, and selective ion transport [7, 11]. Among these mechanisms, Na⁺ ion exclusion from shoot tissues is considered one of the most effective tolerance strategies in cereals [15, 16]. In bread wheat, Na⁺ ion is largely mediated by members of the high-affinity potassium transporter (HKT) family, particularly those encoded by the *Nax* loci introgressed from wild relatives [11]. The *Nax1* gene, corresponding to an HKT1;4-type transporter, functions by retrieving Na⁺ from the xylem sap in roots and leaf sheaths, thereby limiting Na⁺ ion translocation to photosynthetically active tissues [15, 16]. Enhanced expression of *Nax1* has been associated with improved growth and yield stability under saline conditions, highlighting its importance in breeding programs targeting salt-affected environments. Despite advances in identifying salinity tolerance genes, the phenotypic expression of tolerance remains complex and highly genotype-dependent. Early growth traits such as germination rate, survival rate, shoot length, root elongation, and dry matter accumulation are widely used as indicators of salt tolerance in controlled screening systems [9, 17]. Hydroponic systems provide precise control over salinity levels and nutrient composition, allowing reproducible assessment of genotype responses [17]. However, reliance on single traits often fails to capture the multidimensional nature of salinity tolerance.

Multivariate statistical approaches, including PCA and hierarchical clustering, enable integration of multiple physiological parameters to classify genotypes along tolerance–sensitivity gradients [18, 19]. PCA reduces data dimensionality while retaining maximum variance, facilitating identification of key traits contributing to stress resilience. Such integrative approaches are increasingly recommended for screening large germplasm collections under abiotic stress conditions [18, 20]. In parallel, molecular marker systems continue to play a pivotal role in characterizing genetic diversity and supporting marker-assisted selection. SSR markers remain valuable due to their co-dominant inheritance, high polymorphism, and genome-wide distribution [21, 22]. Assessment of polymorphic information content (PIC), gene diversity, and population structure provides insights into the breeding potential of available germplasm. Integration of molecular diversity analysis with phenotypic screening strengthens the identification of elite genotypes possessing both desirable traits and favorable genetic backgrounds [20].

South Asian coastal regions, including large areas of Bangladesh, India, and Pakistan, are increasingly affected by salinity due to tidal intrusion and irrigation-induced salt accumulation [23, 24]. Development of salt-tolerant wheat cultivars adapted to these agroecosystems is therefore essential for maintaining regional food security. While several improved spring wheat genotypes have been released by national breeding programs, comparative physiological and molecular characterization under graded salinity stress remains limited.

Recent studies emphasize that effective salinity tolerance results from coordinated regulation of ion transport, growth maintenance, and stress-responsive gene expression rather than reliance on a single mechanism [7, 12]. In particular, linking *Nax1* expression dynamics with physiological performance under defined salinity gradients can enhance mechanistic understanding and improve the precision of breeding strategies. Therefore, the present study aimed to (i) evaluate growth, biomass, germination, and survival responses of spring wheat genotypes under graded salinity stress; (ii) classify genotypes using multivariate statistical approaches; (iii) assess genetic diversity using SSR and *Nax*-linked markers; and (iv) quantify relative expression of *Nax1* under salinity conditions. By integrating physiological, molecular, and gene expression analyses, this work seeks to advance understanding of salinity tolerance mechanisms in wheat and provide a robust framework for breeding cultivars adapted to salt-affected environments.

## Materials and Methods

### Plant materials

100 genotypes of spring wheat (*Triticum aestivum* L.) were used in the preliminary trials, which included nine wheat varieties (released by the Bangladesh Agricultural Research Institute (BARI) and Bangladesh Wheat and Maize Research Institute (BWMRI)), 52 BAW lines (BAW = Bangladesh Advanced Wheat), 36 BWSN lines (BWSN = Bangladesh Wheat Screening Nursery), and three HZWYT lines (HZWYT = High Zinc Wheat Yield Trial), developed by BWMRI. The pedigree of 100 genotypes was represented in Table S1 [25, 26].

### Development of the hydroponic platform, preparation of Hoogland solutions, and preparation of saline solutions

The development of the hydroponic platform and the preparation of Hoogland solutions and saline solutions were performed as described in File S1.

### Plant materials and experimental design

The preliminary experiments were conducted with 100 genotypes of spring wheat to select suitable genotypes for conducting salinity trials in the hydroponic conditions. Each trial was set up, followed by a randomized complete block design with three replications.

### Measurement of shoot and root length and their dry matter

15 seeds of each genotype were sown on a plate set on a tray filled with tap water. One day after sowing (DAS), to supply nutrients to the seeds/seedlings, the Hoogland solution was also added to the tap water of each tray, and the pH of the solution of the tray was measured with a pH meter to maintain it between 6.0 and 7.0. After 10 DAS, either 0.0 (tap water) or 10, 12, or 14 dS m⁻¹ concentrated saline solution was added to each tank [12]. This tank solution was sent to a tray from where seedlings absorbed saline solution by soft fabric yarn. The saline concentration of tray water was measured by an EC meter. At 20 DAS, 10 seedlings were harvested to measure the morpho-physiological parameters. The shoot and root length was scaled with a scale and averaged. From the next trial conducted by the above-stated method, 10 seedlings were collected, separated into shoots and roots with a knife, packed in brown paper envelopes separately, and kept in the oven where a 70°C temperature was maintained for 72 hours. Shoot and root dry mass was weighed with a 500 mg measuring balance. The results were presented in Table S2.

### Quantification of germination rate

To measure germination rate, the experiments were conducted as per the above-stated methods. A seed is considered germinated when the plumule becomes longer than half the length of the seed and the radicle is equal to or longer than the seed length. The seedlings’ stunted primary roots are considered abnormally germinated [27]. From three DAS, the number of germinated seedlings was counted each day, and it continued up to 12 DAS. The germination rate was obtained by dividing the number of germinated seeds by the total number of seeds set for germination, multiplied by 100 [28].

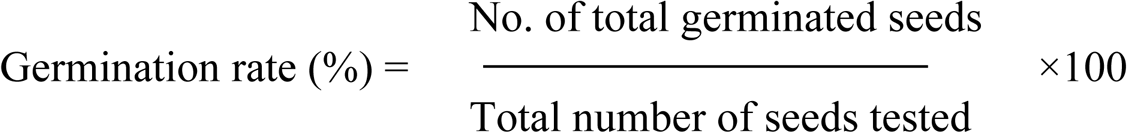

The genotype that exhibited a 50% germination rate was used in the next experiments for further exploration. The experiment was repeated three times for data consistency.

### Quantification of survival rate

To measure the survival rate, the trials were performed using the above-stated methods. Briefly, 15 seeds of each genotype were sown on a plate set on a tray filled with tap water. After 10 DAS, either saline solution or tap water was applied to seedlings (Hasanuzzaman et al., 2020). After five days of saline solution application, the number of seedlings was counted, followed by a one-day interval, and continued up to 25 DAS. At 25 DAS, the survival rate was quantified as the following formula [25, 29]:

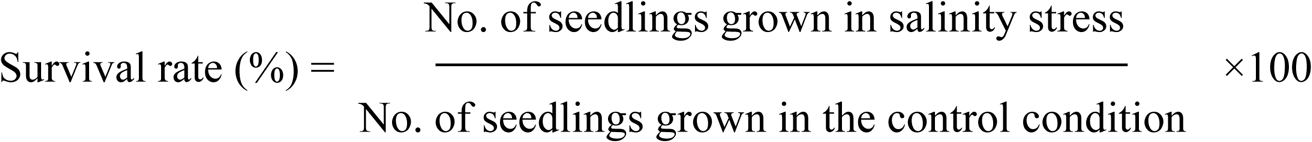

### DNA extraction and molecular marker analysis

Young leaf tissues were collected from 14-day-old seedlings grown under control conditions. Genomic DNA was extracted using the cetyltrimethylammonium bromide method with minor modifications [30]. DNA quality and concentration were assessed using agarose gel electrophoresis and spectrophotometry. A set of SSR markers and *Nax*-linked markers previously reported to be associated with salinity tolerance were selected for amplification (Table S3) [15, 22]. PCR was performed in 20 μL reaction volumes containing genomic DNA, primers, dNTPs, buffer, MgCl₂, and Taq DNA polymerase. Amplified products were resolved on 2-3% agarose gels and visualized under UV light after ethidium bromide staining. Marker bands were scored as binary data (presence = 1, absence = 0). PIC and Nei’s gene diversity (He) were calculated to estimate allelic diversity [21, 31]. Analysis of molecular variance (AMOVA) was performed to partition genetic variation among and within groups. Population structure was further explored using PCA and hierarchical clustering based on marker data (Fig. S1, Fig. S2). Based on the morpho-physiological traits and SSR marker performances of 100 genotypes (Table S2, Fig. S1, Fig. S2), 17 genotypes were selected for the final trials.

### Quantification of physiological traits in the final experiments

17 genotypes were used in the final study. The experiments were performed using the above-stated methods and the protocols established by Zörb et al. (2019), llis & Roberts (1981) and Alam et al. (2022) [14, 29, 32]. The shoot and root length, shoot and root dry matter, total dry matter, germination, and survival rate were recorded as described in the above-stated methods.

### DNA extraction and molecular marker analysis

DNA extraction from leaf and root tissues, PCR, and molecular analysis of the 17 genotypes were performed using the above-stated methods.

### RNA extraction and quantitative RT-PCR analysis

Root and leaf tissues were sampled 48 h after exposure to target salinity levels (0, 10, 12, and 14 dS m⁻¹). Samples were immediately frozen in liquid nitrogen and stored at -80°C until analysis. Total RNA was extracted using a commercial plant RNA isolation kit following the manufacturer’s protocol. RNA integrity was confirmed by agarose gel electrophoresis, and concentration was measured spectrophotometrically. First-strand complementary deoxyribonucleic acid synthesis was performed using reverse transcriptase and oligo (dT) primers. qRT-PCR was conducted using gene-specific primers targeting the sodium transporter gene *Nax1*. Reactions were carried out on a Bio-Rad CFX96 Real-Time™ system SsoAdvanced Universal SYBR Green Supermix™ (Bio-Rad, Hercules, Sweden) following the manufacturer’s protocol. Thermal cycling conditions followed standard protocols for gene expression analysis in wheat (James et al. 2019). Relative gene expression was calculated using the 2⁻ΔΔCt method [33] with actin used as an internal reference gene.

### Statistical analysis

All experiments were conducted hydroponically using a randomized complete block design (RCBD) with three biological replicates, and each experiment was repeated three times to ensure reproducibility. Data were first tested for normality and homogeneity of variance prior to analysis. Differences among treatments and genotypes were evaluated by analysis of variance (ANOVA), followed by Tukey’s honestly significant difference (HSD) test at *P* ≤ 0.05 using IBM SPSS Statistics (version 26.0). Multivariate and population genetic analyses were performed in Python (version 3.14.0). Principal component analysis (PCA) was conducted using standardized (z-score transformed) data to integrate multiple physiological traits. Hierarchical clustering was performed using Ward’s method with Euclidean distance to classify genotypes into salinity tolerance groups. Pearson’s correlation coefficients were calculated to assess relationships between physiological traits and relative *Nax1* gene expression levels, with significance determined at *P* ≤ 0.05. Genetic similarity among wheat genotypes was estimated using Jaccard’s coefficient based on binary marker data. An unweighted pair group method with arithmetic mean (UPGMA) dendrogram was constructed in MEGA (version 12.1) to determine genetic relationships among genotypes under different salinity treatments (0, 10, 12, and 14 dS m⁻¹).

## Results

### Shoot and root elongation were progressively inhibited by increasing salinity

Both shoot length (SL) and root length (RL) decreased significantly with increasing salinity (Fig. 1a-b; 1f-g).

**Fig 1.**
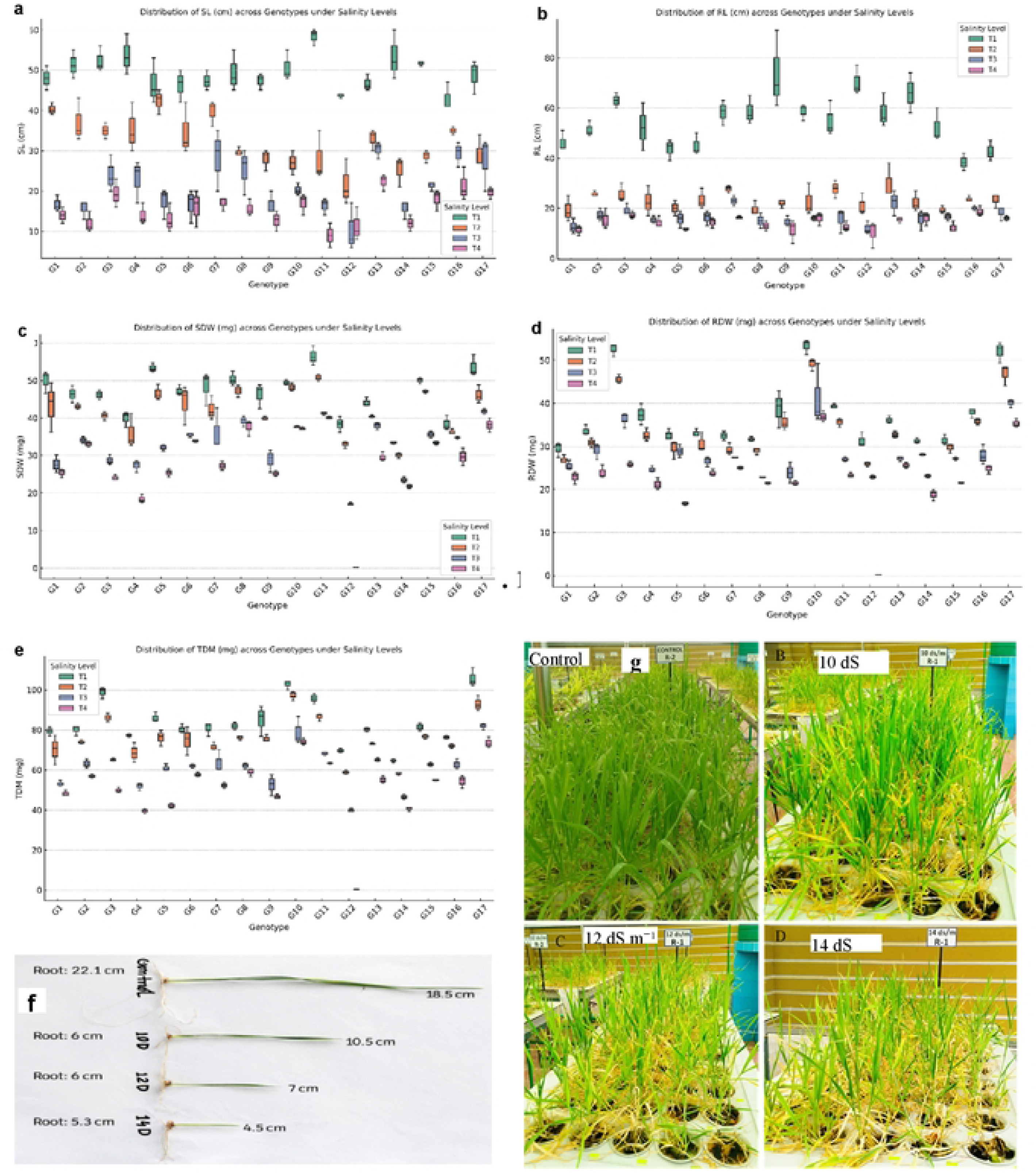
The responses of wheat seedlings to different levels of salinity stress grown in the hydroponic conditions. **a.** Shoot length, **b.** Root length, **c.** Shoot dry weight, **d.** Root dry weight, **e.** Total dry weight, **f.** Scaled shoot and root length, **g.** Phenotypic expressions. The seedlings were grown under controlled conditions. At 10 days after sowing (DAS), different doses of saline solutions (0.0 dS m⁻¹, 10 dS m⁻¹, 12 dS m⁻¹, and 14 dS m⁻¹) were applied to the seedlings. At 20 DAS, the seedlings were harvested, and then shoot and root lengths were scaled. Further, shoots were separated from the root with a sharp knife, packed in brown paper envelopes, kept in an oven at 70°C for 72 hours until constant weight was reached, and finally weighed. Total dry weight was obtained by adding shoot and root weights. The pictures of seedlings treated by different doses of saline solutions were flashed. The **s**hoot and root length were scaled up. Salinity level: T1 = 0.0 dS m⁻¹ (control), T2 = 10 dS m⁻¹, T2 = 12 dS m⁻¹, T2 = 14 dS m⁻¹, Genotypes: G1 = BARI Gom 25, G2 = BARI Gom 30, G3 = BARI Gom 32, G4 = BARI Gom 33, G5 = BWMRI Gom 1, G6 = BWMRI Gom 2, G7 = BWMRI Gom 3, G8 = BWMRI Gom 4, G9 = BWMRI Gom 5, G10 = BAW 1243, G11 = BAW 1286, G12 = BAW 1340, G13 = BAW 1390, G14 = BAW 1422, G15 = BAW 1425, G16 = BWSN 1-14, G17 = BWSN 1-37.

Under control conditions (0.0 dS m⁻¹), genotypes exhibited comparable shoot and root growth. Exposure to 10 dS m⁻¹ resulted in moderate reductions in elongation, whereas 12 and 14 dS m⁻¹ caused substantial inhibition. Shoot elongation declined by approximately 20-25% at 12 dS m⁻¹ and exceeded a 35% reduction at 14 dS m⁻¹ relative to control means. Root elongation was particularly sensitive at higher salinity, with reductions exceeding 40% in sensitive genotypes at 14 dS m⁻¹ (Fig. 1b). Notably, tolerant genotypes retained significantly greater SL and RL compared with sensitive lines under both moderate and high salinity treatments, indicating stronger growth maintenance capacity. The differential response of roots and shoots altered the shoot-to-root length ratio at elevated salinity, reflecting modified biomass allocation patterns under stress conditions.

### Biomass accumulation declined substantially under salinity stress

Salinity significantly reduced shoot dry weight (SDW), root dry weight (RDW), and total dry weight (TDW) (Fig. 1c-e). SDW showed a stronger reduction compared with RDW as salinity intensified. At 14 dS m⁻¹, mean SDW declined by more than 40% relative to control, whereas RDW declined by approximately 30-35%. Total dry weight exhibited a progressive and significant reduction across increasing salinity levels (Fig. 1e). Sensitive genotypes showed drastic biomass loss at 12 and 14 dS m⁻¹, while tolerant lines maintained comparatively higher TDW. Biomass retention percentage (relative to control) clearly differentiated tolerant and sensitive genotypes, with tolerant lines retaining >65% of control biomass at 12 dS m⁻¹ and >50% at 14 dS m⁻¹. These findings indicate that biomass maintenance is a robust physiological indicator of salinity tolerance at the seedling stage.

### Salinity markedly reduced germination and survival rate in a genotype-dependent manner

Salinity significantly affected germination percentage (GP), survival percentage (SP), and all measured growth parameters (P ≤ 0.05) (Fig. 2a-b). GP declined progressively with increasing EC levels (Fig. 2a).

**Fig 2.**
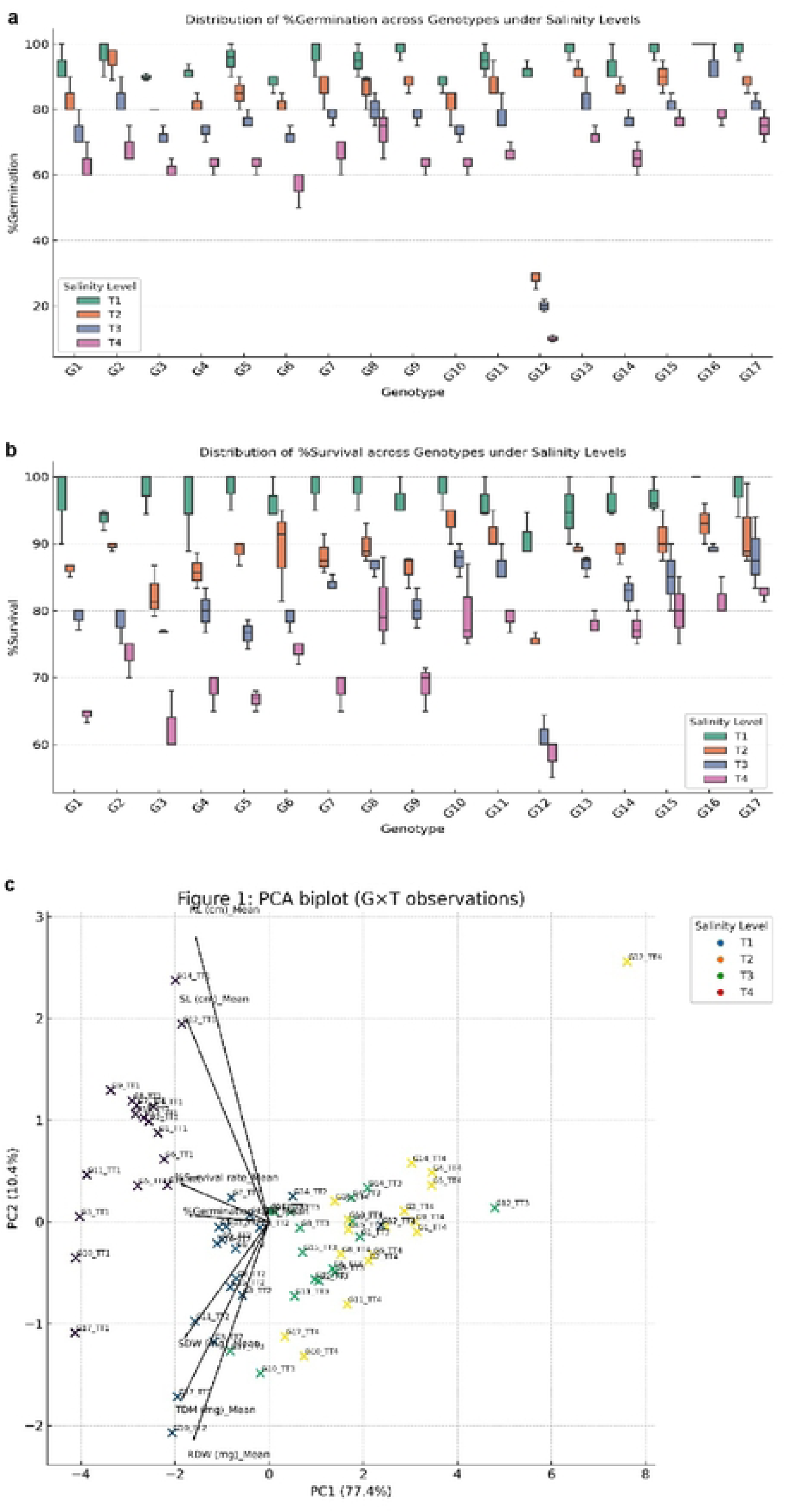
Effects of salinity stress on germination and survival rate and exhibits the principal component analysis (PCA) of physiological traits of wheat seedlings. **a.** Germination rate, **b.** Survival rate, **c.** PCA. 15 seeds were sown in different levels of salinity stress (0.0 dS m⁻¹, 10 dS m⁻¹, 12 dS m⁻¹, and 14 dS m⁻¹). From three days after sowing (DAS), the number of germinated seeds as seedlings was counted, and it continued up to 12 DAS. Then, the germination rate was calculated as described by Cokkizgin (2012). For quantification of survival rate, 15 seeds of each genotype were sown on a tray filled with tap water in the hydroponic conditions. After 10 DAS, salinity stress (above-stated salinity levels) was applied to seedlings except for the control. After five days of stress application, the number of seedlings was counted, followed by a one-day interval, and continued up to 25 DAS, and the survival rate was quantified as described by Ellis & Roberts (1981) and Alam et al. (2022). PCA biplot based on shoot length (SL), root length (RL), shoot dry weight (SDW), root dry weight (RDW), total dry weight (TDW), germination, and survival rate of wheat genotypes under salinity stress. The first two principal components explain the majority of total variation and separate genotypes along a tolerance-sensitivity gradient. Salinity level: T1 = 0.0 dS m⁻¹ (Control), T2 = 10 dS m⁻¹, T2 = 12 dS m⁻¹, T2 = 14 dS m⁻¹, T = Treatment; Genotypes: G1 = BARI Gom 25, G2 = BARI Gom 30, G3 = BARI Gom 32, G4 = BARI Gom 33, G5 = BWMRI Gom 1, G6 = BWMRI Gom 2, G7 = BWMRI Gom 3, G8 = BWMRI Gom 4, G9 = BWMRI Gom 5, G10 = BAW 1243, G11 = BAW 1286, G12 = BAW 1340, G13 = BAW 1390, G14 = BAW 1422, G15 = BAW 1425, G16 = BWSN 1-14, G17 = BWSN 1-37.

Under 10 dS m⁻¹, the mean reduction in GP across genotypes was moderate, whereas 12 and 14 dS m⁻¹ caused pronounced inhibition. At 14 dS m⁻¹, average germination decreased by approximately one-third compared with control conditions. However, substantial genotypic variation was observed. Several genotypes maintained relatively high germination (>75%) even at 12 dS m⁻¹, whereas sensitive lines showed sharp declines. SP was more strongly affected than germination (Fig. 2b). At 12 and 14 dS m⁻¹, SP declined markedly in sensitive genotypes, with some lines exhibiting less than 50% survival at the highest salinity level. In contrast, tolerant genotypes maintained comparatively higher SP (>65%) under 12 dS m⁻¹ and retained appreciable viability at 14 dS m⁻¹. The genotype × salinity interaction was significant, indicating differential tolerance among lines.

### Principal component analysis discriminated between tolerant and sensitive genotypes

To integrate multiple physiological traits, PCA was conducted using standardized data (Fig. 2c). The first two principal components (PC1 and PC2) together explained 42.09% of the total variation. PC1 accounted for the largest proportion of variance and was strongly associated with TDW, SDW, SP, and SL. PC2 was primarily associated with RL and RDW. The PCA biplot clearly separated genotypes along a tolerance–sensitivity gradient. Tolerant genotypes clustered on the positive side of PC1, characterized by higher biomass retention and survival under salinity stress. Sensitive genotypes grouped on the negative side of PC1, reflecting pronounced reductions in growth parameters. The distribution pattern indicated that biomass and survival contributed most significantly to overall salinity tolerance variation.

### Hierarchical clustering confirmed distinct tolerance groups

Hierarchical clustering using Ward’s method based on physiological traits further classified genotypes into distinct clusters (Fig. 3).

**Fig 3.**
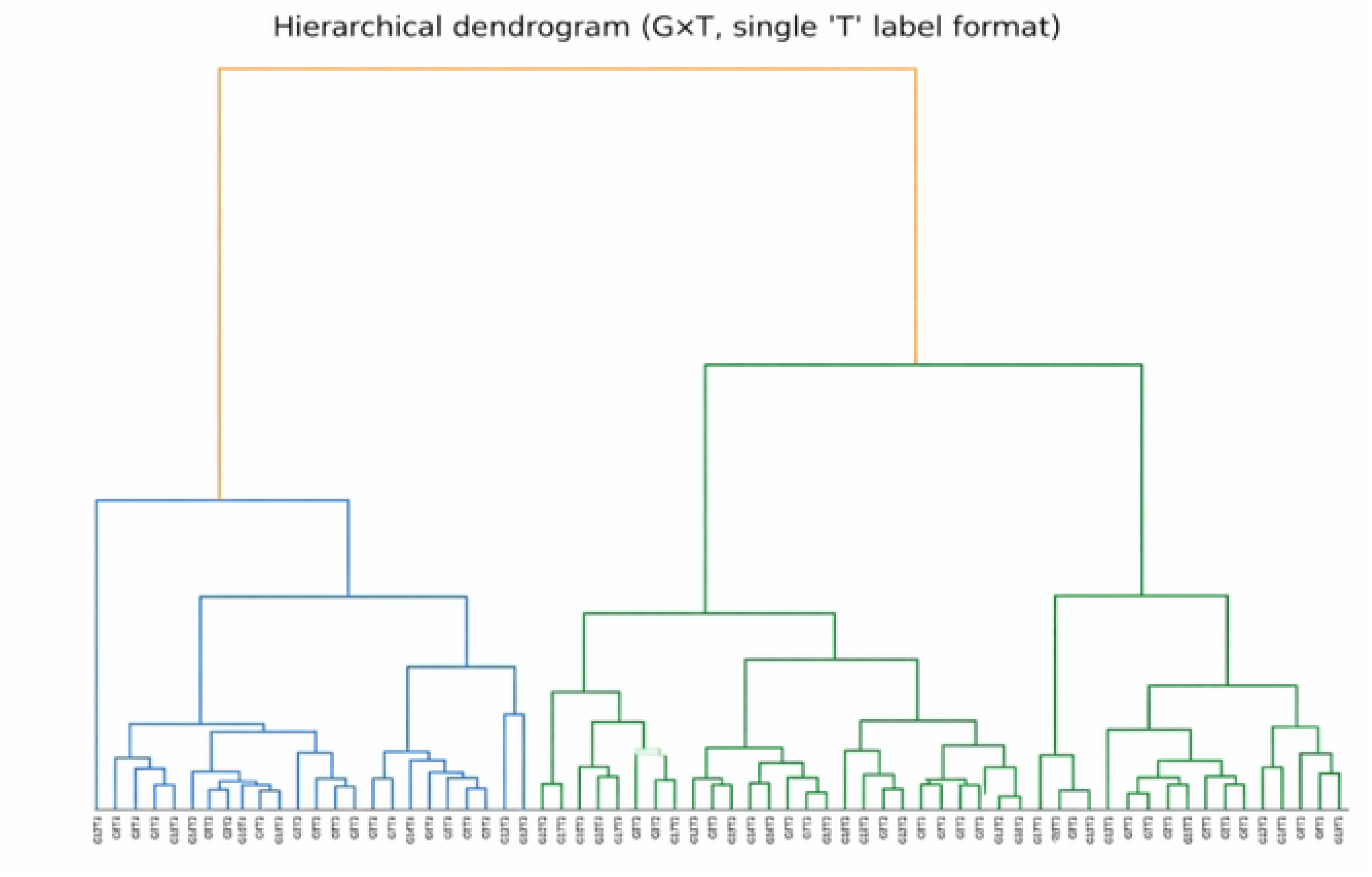
Hierarchical clustering of wheat genotypes based on physiological performances. Hierarchical clustering dendrogram constructed using standardized physiological and biomass traits under salinity stress. Genotypes were grouped into tolerant and sensitive clusters, supporting multivariate classification of salinity responses. Salinity level: T1 = 0.0 dS m⁻¹ (Control), T2 = 10 dS m⁻¹, T2 = 12 dS m⁻¹, T2 = 14 dS m⁻¹, T = Treatment; Genotypes: G1 = BARI Gom 25, G2 = BARI Gom 30, G3 = BARI Gom 32, G4 = BARI Gom 33, G5 = BWMRI Gom 1, G6 = BWMRI Gom 2, G7 = BWMRI Gom 3, G8 = BWMRI Gom 4, G9 = BWMRI Gom 5, G10 = BAW 1243, G11 = BAW 1286, G12 = BAW 1340, G13 = BAW 1390, G14 = BAW 1422, G15 = BAW 1425, G16 = BWSN 1-14, G17 = BWSN 1-37.

Two major groups were identified: Cluster I comprised genotypes exhibiting superior growth and biomass retention under 12-14 dS m⁻¹, whereas Cluster II included sensitive genotypes with substantial reductions in survival and dry matter accumulation. Sub-clustering within the tolerant group suggested varying degrees of tolerance, indicating quantitative differences among genotypes. The clustering pattern corresponded closely with PCA classification, reinforcing the reliability of multivariate approaches for salinity tolerance screening.

### Genetic diversity and phylogenetic relationships among wheat **g**enotypes revealed by Neighbor-joining Analysis

The circular Neighbor-Joining (NJ) phylogenetic tree constructed from molecular marker data revealed clear genetic differentiation among the wheat genotypes (Fig. 4).

**Fig 4.**
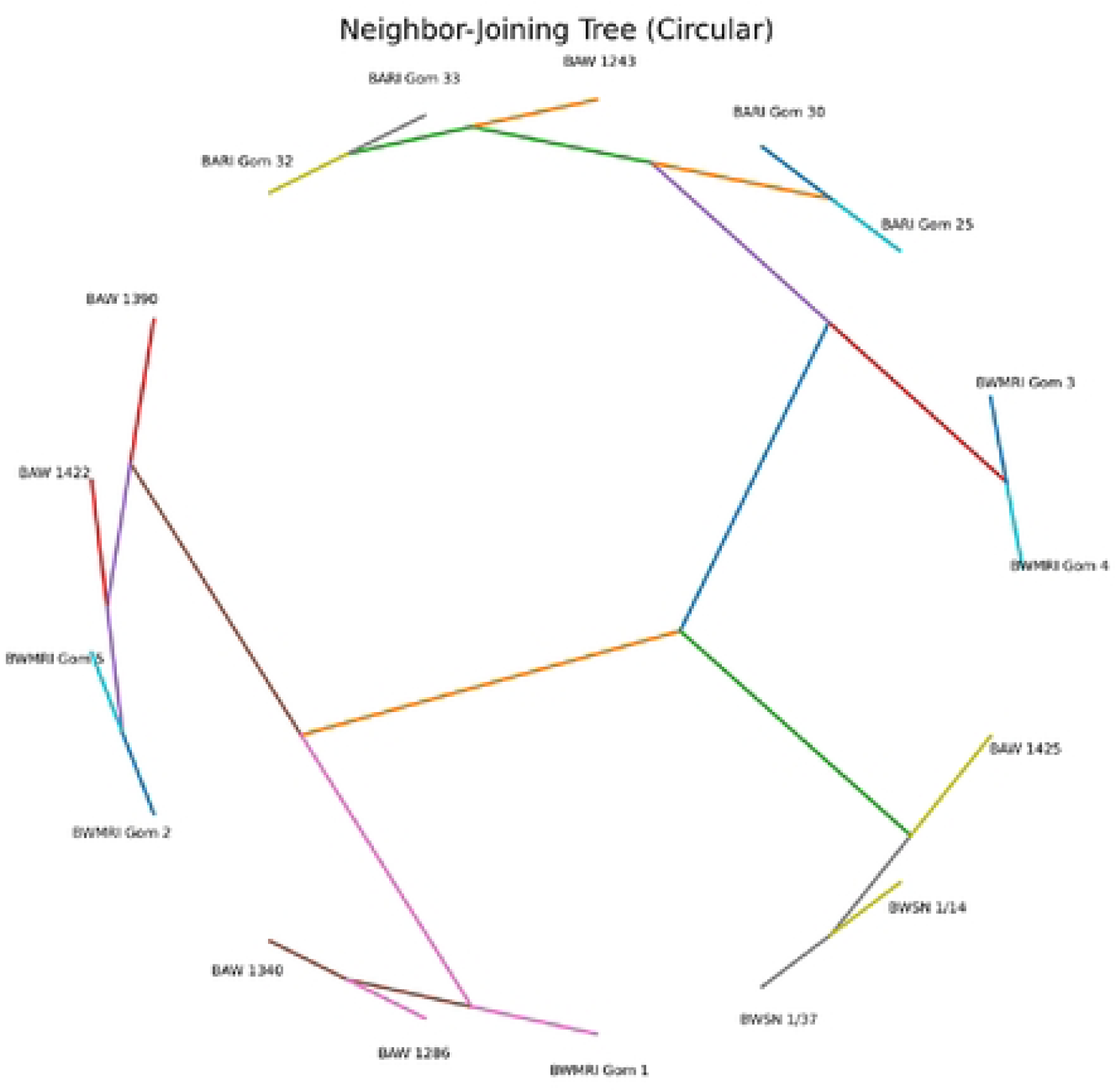
Circular Neighbor-Joining (NJ) phylogenetic tree depicting the genetic relationships among wheat genotypes based on molecular marker data. The tree was constructed using a pairwise genetic distance matrix derived from marker polymorphism profiles. Each terminal node represents an individual genotype, and branch lengths correspond to genetic distances among genotypes. The circular layout facilitates clear visualization of clustering patterns, revealing genetic similarity and divergence within the population, which may be associated with stress-adaptive traits.

The clustering pattern separated the genotypes into several distinct groups, indicating substantial genetic variability within the studied population. Genotypes belonging to similar pedigree backgrounds or breeding programs tended to cluster together, reflecting shared ancestry and genetic similarity. Branch lengths varied considerably among genotypes, suggesting differences in genetic distance and divergence levels. Some genotypes formed tight sub-clusters with short branch lengths, indicating close genetic relationships, whereas others appeared as distinct lineages with longer branches, reflecting higher genetic divergence. The overall topology demonstrates moderate-to-high polymorphism across the molecular markers used, confirming the effectiveness of the marker system in discriminating among wheat genotypes. The clustering pattern suggests the presence of genetically diverse groups that could serve as potential parental lines for breeding programs targeting stress tolerance and yield stability.

### SSR and *Nax*-linked markers revealed moderate genetic diversity

Molecular analysis using SSR and *Nax*-linked markers generated clear and reproducible banding patterns. Binary scoring of amplified fragments indicated moderate polymorphism across loci (Fig. 5a).

**Fig 5.**
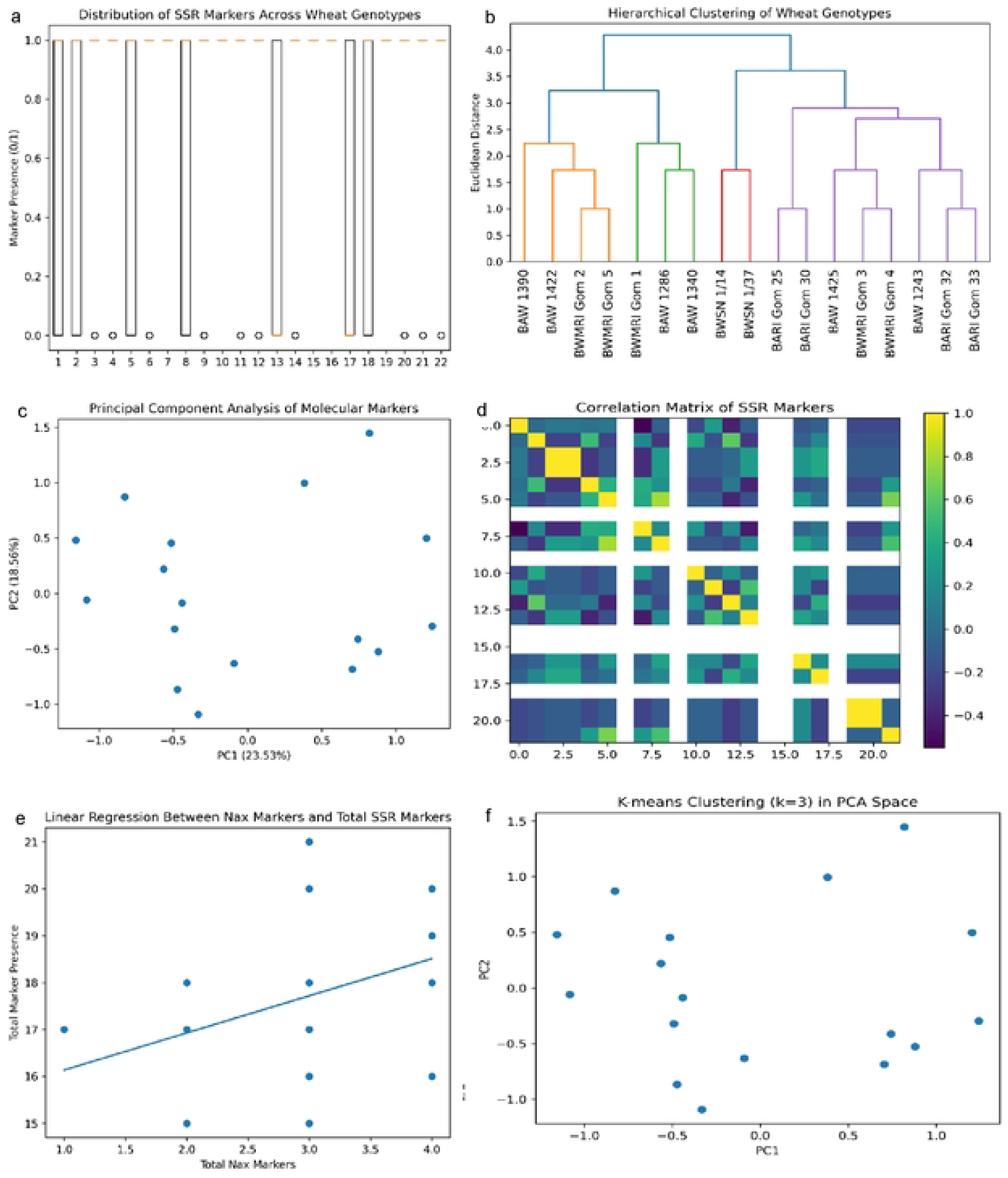
Multivariate and population genetic analysis of wheat genotypes using SSR. **a**. Boxplot illustrating distribution of binary marker scores (1/0), indicating moderate polymorphism across loci, **b.** Ward’s hierarchical clustering dendrogram based on Euclidean genetic distance, revealing three principal genetic groups, **c.** Principal component analysis (PCA) showing genetic differentiation; PC1 and PC2 accounted for 23.53% and 18.56% of total variance (42.09% cumulative), respectively. **d.** Pearson correlation matrix demonstrating predominantly positive inter-locus associations, suggesting possible linkage or shared chromosomal regions. **e.** Linear regression analysis indicates a positive relationship between total *Nax*-linked alleles and overall marker presence. **f.** K-means clustering (k = 3) projected onto PCA space, confirming population structure. Mean polymorphic information content (PIC) and Nei’s gene diversity (He) indicated moderate allelic variation, while AMOVA revealed minimal among-group differentiation (ΦST ≈ 0.0006), suggesting that most genetic variance resides within genotypes rather than between clusters.

PIC values suggested moderate allelic diversity among the studied genotypes. Ward’s hierarchical clustering based on marker-derived genetic distance grouped genotypes into three principal genetic clusters (Fig. 5b). PCA of marker data further supported genetic differentiation, with PC1 and PC2 explaining 23.53% and 18.56% of total molecular variation, respectively (42.09% cumulative; Fig. 5c). Analysis of molecular variance (AMOVA) indicated minimal among-group differentiation (Φ_ST ≈ 0.0006), demonstrating that most genetic variation resided within genotypes rather than between clusters. Pearson correlation analysis of marker loci revealed predominantly positive inter-locus associations (Fig. 5d). K-means clustering projected onto PCA space confirmed the presence of three genetic groups (Fig. 5e-f). Importantly, tolerant and sensitive genotypes were distributed across different molecular clusters, suggesting that physiological tolerance was not strictly confined to a single genetic background.

### *Nax1* expression was strongly induced in tolerant genotypes under severe salinity

Relative expression analysis demonstrated significant induction of *Nax1* in response to salinity (Fig. 6).

**Fig 6.**
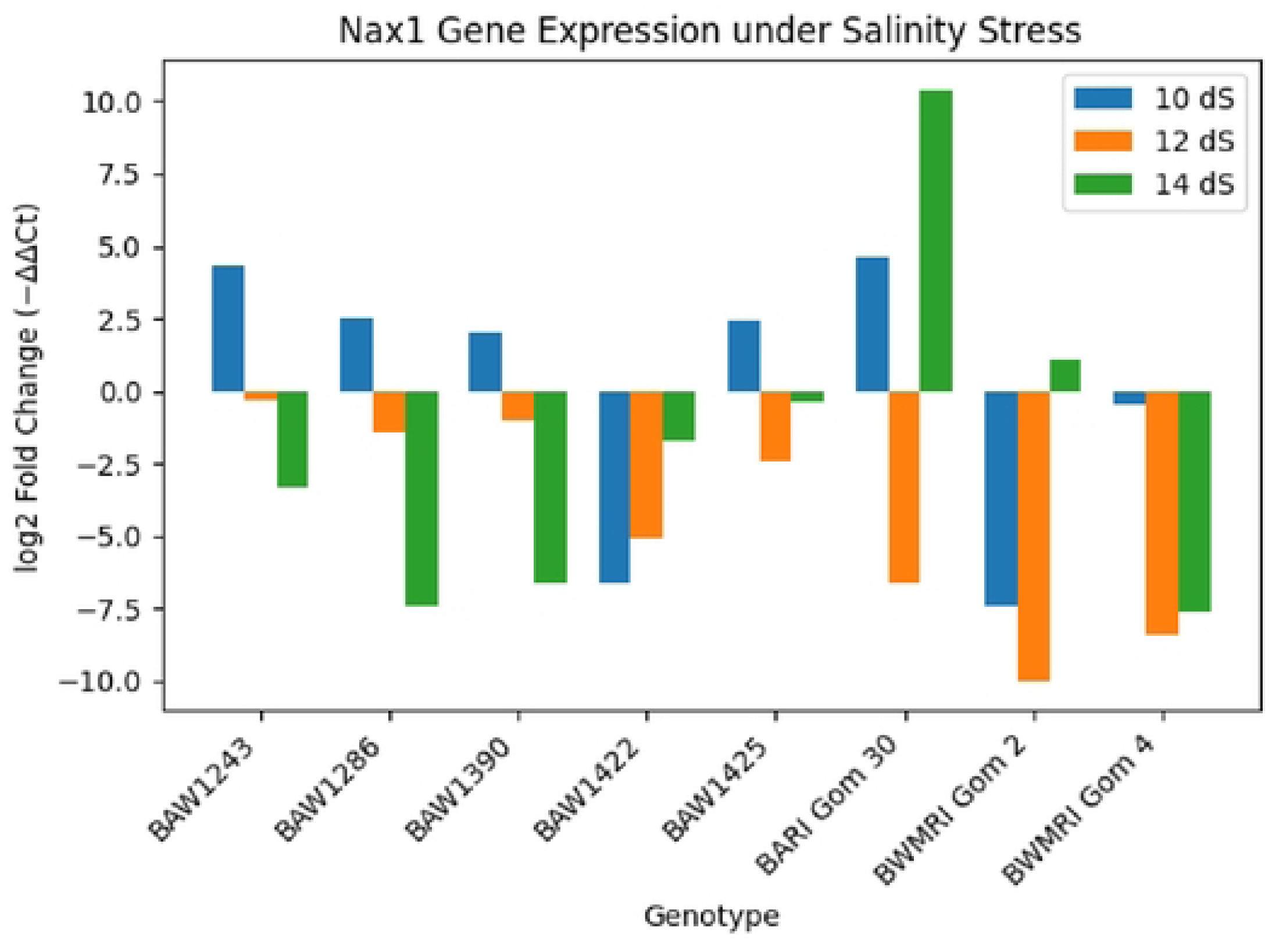
Relative expression of the sodium exclusion gene *Nax1* under salinity stress. Relative expression levels of *Nax1* in wheat genotypes exposed to salinity stress (10, 12, and 14 dS m⁻¹), expressed as log₂ fold change relative to control conditions. Tolerant genotypes exhibit stronger and more sustained *Nax1* induction under increasing salinity.

At 10 dS m⁻¹, moderate upregulation was observed across most genotypes. However, at 12 and 14 dS m⁻¹, tolerant genotypes exhibited pronounced induction (3-6-fold increase relative to control), whereas sensitive genotypes showed comparatively limited upregulation. Expression levels increased progressively with salinity intensity in tolerant lines, indicating stress-responsive regulation. In contrast, sensitive genotypes displayed either weak induction or inconsistent expression patterns under high salinity. Correlation analysis revealed a positive association between relative *Nax1* expression and total dry weight retention under 12-14 dS m⁻¹. Genotypes with higher *Nax1* induction consistently maintained greater biomass and survival, suggesting a functional linkage between sodium exclusion and growth maintenance.

### Integrated physiological and molecular framework

A schematic integration of physiological performance and molecular regulation was presented in Fig. 7.

**Fig 7.**
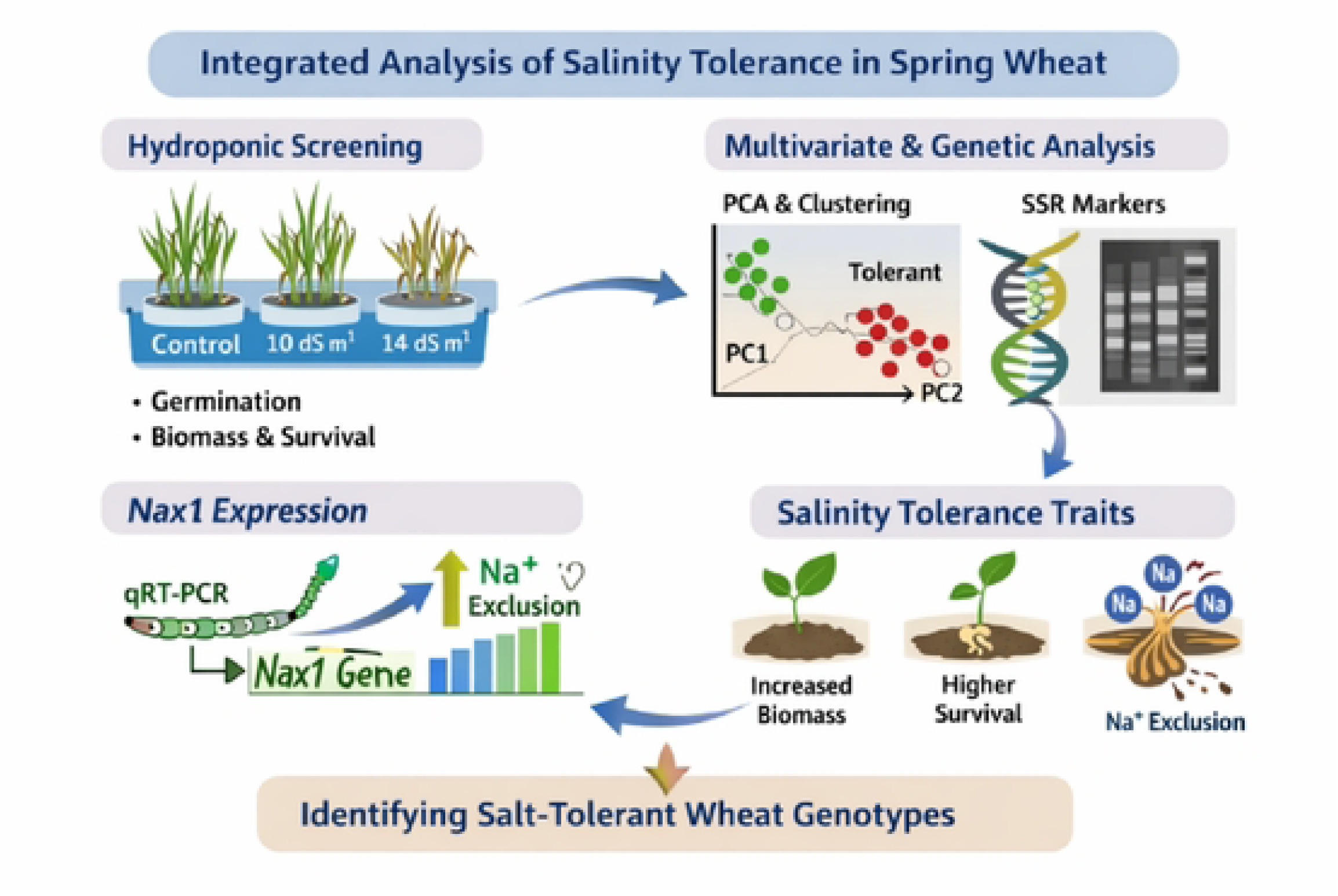
Schematic overview of the integrated approach for evaluating salinity tolerance in spring wheat. Hydroponic screening was conducted under 0.0 (control), 10 dS m⁻¹, and 14 dS m⁻¹ salinity levels to assess germination, biomass, and survival. Multivariate analyses, including principal component analysis (PCA) and clustering, were applied to classify genotypes based on tolerance performance, and genetic diversity was examined using SSR markers. Relative expression of the *Nax1* gene was quantified by qRT-PCR to evaluate its role in Na⁺ exclusion. Key salinity tolerance traits—enhanced biomass, improved survival, and efficient Na⁺ exclusion-were integrated to identify salt-tolerant wheat genotypes.

Salinity-induced reductions in germination, elongation, and biomass were mitigated in tolerant genotypes through sustained *Nax1*-mediated sodium exclusion. Multivariate trait analysis, marker diversity assessment, and gene expression profiling collectively provided a coherent classification of genotypes along a tolerance gradient. Overall, the results demonstrate significant genotype-dependent variation in salinity tolerance and highlight biomass retention and *Nax1* expression as key determinants of resilience under saline conditions.

## Discussion

Recent molecular investigations highlight the regulatory role of Na⁺ transporters, particularly HKT-mediated sodium retrieval and tissue-specific ion partitioning, in enhancing salt tolerance [34-38]. These findings support the physiological and molecular patterns observed in the present study. Salinity stress significantly impaired germination, survival, shoot and root growth, and biomass accumulation in spring wheat, confirming the sensitivity of early developmental stages to osmotic and ionic stress (Fig. 1-2). The progressive decline in growth parameters with increasing EC reflects the biphasic nature of salinity stress: an initial osmotic phase restricting water uptake followed by ionic toxicity resulting from excessive Na⁺ accumulation [8-11]. The stronger reduction in survival compared with germination suggests that prolonged exposure to saline conditions exacerbates ionic imbalance and metabolic disruption, consistent with previous reports in wheat and other cereals [9, 14].

Biomass retention emerged as the most reliable physiological indicator of tolerance among the studied genotypes. Sensitive lines exhibited substantial reductions in shoot dry weight and total dry matter at 12-14 dS m⁻¹, whereas tolerant genotypes maintained comparatively higher biomass (Fig. 1c-e). Maintenance of biomass under salinity is generally associated with effective ion homeostasis, osmotic adjustment, and sustained photosynthetic activity [7, 12]. Excess Na⁺ disrupts K⁺ uptake and enzyme activation, leading to impaired carbon assimilation and growth inhibition [14, 39]. Therefore, the ability of tolerant genotypes to sustain dry matter accumulation suggests improved regulation of ion transport and cellular osmotic balance. Root growth was particularly sensitive at higher salinity levels, reflecting the direct exposure of roots to saline solution and the importance of root-based ion transport processes. Reduced root elongation in sensitive genotypes may indicate impaired cell expansion and membrane integrity due to Na⁺ toxicity (Fig. 1b, d-e) [10]. Conversely, tolerant genotypes maintained relatively greater root length and root dry weight, which may facilitate improved water uptake and nutrient acquisition under stress. Enhanced root performance under salinity has been linked to improved Na⁺ retrieval from the xylem and more efficient exclusion mechanisms [15].

The application of multivariate statistical approaches provided further insight into the complex nature of salinity tolerance. PCA revealed that biomass-related traits and survival contributed most strongly to overall variation, reinforcing their importance as integrative indicators of tolerance. The clear separation of genotypes along a tolerance-sensitivity gradient demonstrates the value of PCA for reducing data dimensionality while retaining biologically meaningful variation (Fig. 2c) [18, 19]. Hierarchical clustering corroborated PCA results, grouping genotypes into tolerant and sensitive clusters (Fig. 3). Such integrative classification approaches are increasingly recommended for screening large germplasm sets under abiotic stress conditions (Bhatta et al. 2018).

The NJ-based clustering revealed substantial genetic diversity among the wheat genotypes, highlighting valuable variation for stress-resilient breeding. Genetic diversity provides the foundation for selection and adaptation in crop improvement programs [5]. The clear grouping pattern indicates that the molecular markers effectively captured genome-wide polymorphism, enabling reliable discrimination among genotypes. The Neighbor-Joining method is widely applied for assessing genetic relationships from distance matrices [40] and its use in wheat has proven effective for evaluating population structure with molecular markers [41]. Genotypes forming tight clusters likely share common breeding backgrounds, whereas those on longer branches may possess unique alleles linked to adaptive stress responses. Genetically distant parents are often associated with greater heterosis and improved stress resilience [42]. Therefore, the divergent genotypes identified here represent promising candidates for crossing programs. Integrating molecular diversity with stress-related phenotyping strengthens the selection of elite germplasm for climate-resilient wheat breeding [43].

Molecular marker analysis revealed moderate genetic polymorphism among the tested genotypes, indicating the presence of exploitable diversity despite shared breeding histories (Fig. 5). SSR markers remain effective tools for assessing allelic variation and population structure in wheat [21, 41]. The low among-group differentiation observed in AMOVA suggests that most genetic variation resides within genotypes rather than between predefined tolerance groups. This pattern likely reflects the common genetic background of modern cultivars derived from related breeding programs. Importantly, tolerant and sensitive genotypes were distributed across different molecular clusters, implying that salinity tolerance is a quantitative trait influenced by multiple loci rather than confined to a single lineage (Fig. 5).

A central finding of this study is the pronounced induction of *Nax1* expression in tolerant genotypes under severe salinity (12-14 dS m⁻¹) (Fig. 6). The *Nax1* locus encodes an HKT1;4-type sodium transporter responsible for retrieving Na⁺ from the xylem, thereby limiting its accumulation in leaf blades [15, 16]. Enhanced *Nax1* expression in tolerant genotypes suggests activation of a robust Na⁺ exclusion mechanism that protects photosynthetically active tissues from ionic toxicity. The positive correlation between *Nax1* expression and biomass retention supports a functional linkage between molecular regulation and physiological performance (Fig. 7).

Previous studies have demonstrated that introgression of *Nax* loci into bread wheat significantly reduces leaf Na⁺ concentration and improves yield stability under saline field conditions [11]. Our results extend these findings by demonstrating that endogenous variation in *Nax1* expression among spring wheat genotypes contributes to differential tolerance under controlled hydroponic conditions (Fig. 6). The stress-responsive upregulation of *Nax1* observed here aligns with evidence that HKT transporter expression is dynamically regulated in response to Na⁺ exposure [7, 16]. Salinity tolerance is inherently multifactorial, involving not only Na^+^ exclusion but also vacuolar sequestration, osmolyte synthesis, and antioxidant defense [12]. While this study focused primarily on *Nax1*-mediated exclusion, the observed variation in growth performance likely reflects coordinated action of multiple pathways (Fig. 7). Future studies incorporating ion concentration measurements, transcriptomic profiling, and antioxidant enzyme analysis would provide a more comprehensive understanding of the regulatory networks underlying tolerance. The integration of physiological screening, multivariate analysis, molecular diversity assessment, and gene expression profiling represents a robust framework for identifying salt-tolerant wheat genotypes (Fig. 7). By linking biomass maintenance with *Nax1* induction, this study provides mechanistic insight into how Na^+^ exclusion contributes to growth resilience under saline conditions. Such integrative approaches are essential for accelerating marker-assisted breeding and improving wheat adaptation to salt-affected agroecosystems, particularly in regions where soil salinization continues to intensify.

Overall, our findings reinforce the central role of Na^+^ exclusion in wheat salinity tolerance and demonstrate that variation in *Nax1* expression significantly contributes to genotype-dependent resilience. The combination of physiological and molecular analyses enhances understanding of the complexity of salinity adaptation and supports targeted improvement of wheat for saline environments.

## Conclusion

This study demonstrates substantial genotype-dependent variation in salinity tolerance among spring wheat at the seedling stage. Increasing salinity (10-14 dS m⁻¹) significantly reduced GP, SP, SL, RL, SDW, RDW, and biomass accumulation. However, tolerant genotypes maintained higher biomass retention and SP under severe salinity, highlighting their superior adaptive capacity. Among the evaluated physiological parameters, TDW and SP proved to be the most reliable indicators for discriminating between tolerant and sensitive genotypes. Multivariate analyses effectively integrated multiple traits and clearly separated genotypes along a tolerance gradient, emphasizing the utility of PCA and hierarchical clustering in stress screening programs. The phylogenetic analysis revealed significant genetic diversity among the 17 wheat genotypes. The dendrogram provides a valuable genetic framework for selecting parental lines in breeding programs aimed at improving salinity tolerance. The circular Neighbor-Joining analysis revealed clear genetic structuring and substantial diversity among the studied wheat genotypes. The distinct clustering pattern confirms the effectiveness of the molecular markers in detecting genome-wide polymorphism and differentiating closely related materials. Genotypes located on longer branches represent genetically divergent resources that may harbor unique alleles associated with stress adaptation, while closely clustered genotypes likely share common breeding backgrounds.

A key finding of this study is the strong induction of *Nax1* expression in tolerant genotypes under high salinity. The positive association between *Nax1* upregulation and biomass maintenance supports the central role of Na^+^ exclusion in mitigating ionic toxicity. These results provide functional evidence linking molecular regulation of Na⁺ transport with improved growth performance under saline conditions. Collectively, the integration of physiological, multivariate, and molecular analyses advances our understanding of salinity adaptation in wheat (Fig. 11). The identified tolerant genotypes and the demonstrated relevance of *Nax1* expression offer valuable resources for marker-assisted selection and breeding programs aimed at improving wheat productivity in salt-affected environments.

## Abbreviations

AMOVA: Analysis of molecular variance
BARI: Bangladesh Agricultural Research Institute
BAW: Bangladesh Advanced Wheat
BWMRI: Bangladesh Wheat and Maize Research Institute
DAS: Day after sowing
DNA: Deoxyribonucleic acid
GP: germination percentage
PCA: Principal component analysis
PCR: Polymerase chain reaction
PIC: Polymorphic information content
qRT-PCR: Quantitative real-time polymerase chain reaction
RDW: Root dry weight
RL: Root length
RNA: Ribonucleic acid
SDW: Shoot dry weight
SL: Shoot length
SP: Survival percentage
SSR: Simple Sequence Repeat
TDW: Total dry weight.

## Authors’ contributions

MH and MAKA conceptualized and designed the experiments. MHH investigated the experiments and curated the data. MNA performed formal analysis, guided methodology, and administered the project. MH and MAKA supervised the experiments. MNA visualized all figures. MNA and MHH wrote the original draft. MAKA and MNA validated data. MH, MAKA, and MNA revised, edited, and improved the manuscript.

## Acknowledgements

We thank Professor Dr. Md. Abu Sayed and Md. Tanvir Abedin (research assistant) for instrument operation guidance (Department of Biochemistry and Molecular Biology, Hajee Mohammad Danesh Science & Technology University, Dinajpur, Bangladesh). Heartiest gratefulness to Dr. Zaherul Islam, Principal Scientific Officer, Wheat Breeding Division, Bangladesh Wheat and Maize Research Institute, Dinajpur, Bangladesh, for his technical support.

## Supplementary information

File S1: Hydroponic platform, Hoogland solutions, saline solution.

Fig S1: Figures of PCR of 100 genotypes were performed with molecular markers.

Fig S2: Molecular diversity and genetic relationships of 100 wheat genotypes.

Table S1: Pedigree of 100 Genotypes.

Table S2: Physiological performance of 100 genotypes.

Table S3: Nax genes and salinity-related markers used for selecting salinity-tolerant wheat genotypes.

Table S4: The gene was used for qRT-PCR.

